# Optimizing Intermittent Theta-Burst Stimulation: The Effects of Pulse Parameters on Corticospinal Excitability

**DOI:** 10.64898/2026.10.05.756785

**Authors:** Béatrice Proulx, Seyede Anis Hasani, Ann-Jessica Fréchette, Raphaël Hamel, Sara Tremblay, Samantha Côté, Jean-François Lepage

**Affiliations:** Department of Psychology, Faculty of Social Sciences, Bishop’s University, Sherbrooke, Canada; Center for Molecular and Behavioral Neurosciences, Rutgers University, Newark, United States of America; Department of Pediatrics, Faculty of Medicine and Health Sciences, Université de Sherbrooke, Sherbrooke, Canada; Department of Medical Imagery Sciences and Radiations, Faculty of Medicine and Health Sciences, Université de Sherbrooke, Sherbrooke, Canada; School of Sport, Exercise, and Rehabilitation Sciences, University of Birmingham, United Kingdom; Department of Psychoeducation and Psychology, Université du Québec en Outaouais, Gatineau, Canada

**Keywords:** Transcranial magnetic stimulation, neurostimulation, iTBS, plasticity, LTP, GABA, major depressive disorder

## Abstract

**Objective:** Intermittent theta burst stimulation (iTBS) is a repetitive transcranial magnetic stimulation (rTMS) protocol used in clinical settings to alleviate symptoms of psychiatric disorders. Although the pulse parameters currently used in iTBS have remained unchanged since the first treatments, recent controllable TMS devices allow for the personalization of pulse shape and duration, opening the possibility of testing the efficacy of other pulse parameters in modulating corticospinal excitability (CSE).

**Methods:** In a repeated-measures design, changes in CSE were induced by the administration of iTBS over the left primary motor cortex under three different conditions: long bidirectional pulse (L-Bi), long quasi-unidirectional pulse (L-QU), and a control short quasi-unidirectional pulse (S-QU). We assessed TMS-induced motor evoked potentials before and up to 70 minutes after iTBS. We hypothesized that L-QU pulses would induce greater facilitation than other conditions.

**Results:** A mixed linear model showed that all conditions induced increased MEPs compared to baseline. However, the L-Bi pulse led to a larger and more sustained MEP increase than both unidirectional pulses, while the S-QU pulse appeared less effective in modulating CSE. We observed large within- and between-subject variability in responsiveness to the iTBS protocols.

**Conclusion:** These results suggest that L-Bi pulses in iTBS protocols are more potent than unidirectional pulses in inducing CSE changes in the cortex of healthy adults. The individual variability in response to iTBS and the potential clinical relevance of these findings are discussed.

## Introduction

Intermittent theta-burst stimulation (iTBS) is a form of repetitive transcranial magnetic stimulation (rTMS) that can modulate cortical excitability. iTBS is currently used in the treatment of several neuropsychiatric disorders, including treatment-resistant major depressive disorder (MDD), in which it is applied to key frontal regions (Chaves et al., 2024). However, about half of patients undergoing iTBS for treatment-resistant depression do not respond positively (Blumberger et al., 2018), highlighting the need to optimize stimulation parameters. One promising avenue is controllable TMS (cTMS) devices, which allow pulse shape and duration to be tailored to each individual, potentially strengthening the long-term potentiation (LTP)-like effects that underlie iTBS’s impact on excitability (Huang et al., 2005).

Until recently, TMS devices could only deliver fixed bidirectional pulses in iTBS protocols, a constraint of their capacitor-discharge design (Peterchev et al., 2014). Controllable TMS (cTMS) technology overcomes this by enabling the manipulation of pulse shape, current direction, and phase asymmetry, which compelled researchers to consider novel dimensions of the TMS pulse, such as the M-ratio (Agboada et al., 2026). The M-ratio quantifies the amplitude of the pulse’s second phase relative to its first phase and therefore provides a measure of pulse asymmetry. Different M-ratios modify the temporal and directional characteristics of the induced electric field, allowing the recruitment of specific neural elements (Halawa et al., 2019). Low ratios (< 0.5) yield asymmetric, unidirectional-like waveforms, while higher ratios (≥ 0.5) yield more symmetric, bidirectional waveforms (Sommer et al., 2018).

Although bidirectional pulses remain standard in rTMS research (Sommer et al., 2018), recent studies support greater effectiveness of unidirectional, low-M-ratio pulses to modulate CSE than bidirectional pulses (Agboada et al., 2026). Namely, using 1 Hz rTMS, unidirectional waveforms elicit stronger inhibitory aftereffects and lower motor thresholds than bidirectional pulses in the motor cortex (Goetz et al., 2016; Halawa et al., 2019). However, single-pulse studies support that bidirectional pulses induce lower motor thresholds and higher MEP amplitudes (Sommer et al., 2006; Delvendahl et al., 2014). Overall, no consensus has been reached on which pulse shape most efficiently modulates cortical excitability (Agboada et al., 2026).

Pulse duration critically shapes iTBS’s excitability effects. Evidence consistently shows that longer pulses more effectively recruit cortical neurons and produce stronger aftereffects; they require lower stimulator intensities to elicit motor responses (Peterchev et al., 2013), are associated with lower resting motor thresholds, shorter MEP latencies and higher MEP amplitudes (Halawa et al., 2019; 2021). Even in 1 Hz rTMS where inhibiton is expected, sending longer pulses reversed the directionality of aftereffects, with inhibition becoming facilitation (Halawa et al., 2019). This likely reflects more effective depolarization of neuronal populations with longer membrane time constants, such as interneurons and dendritic compartments, that require a sustained input to reach threshold and are important for synaptic plasticity (Stern et al., 2015; D’Ostilio et al., 2016). Yet it remains unclear whether pulse duration similarly conditions the effects of bidirectional versus unidirectional pulses on excitability (Agboada et al., 2026).

Despite iTBS’s clinical relevance, few studies have systematically examined how pulse shape, duration, and M-Ratio jointly influence its excitability effects (Agboada et al., 2026); most optimization work has instead focused on single-pulse (Casula et al., 2018; Delvendahl et al., 2014) or low-frequency rTMS (Goetz et al., 2016; Halawa et al., 2019; 2021). To address this gap, in a fully within-subject design, we measured corticospinal excitability (CSE) using single TMS pulses over M1 before and up to 70 minutes after an iTBS intervention that delivered long bidirectional (L-Bi), long quasi-unidirectional (L-QU), or control short quasi-unidirectional pulses (S-QU). In this study, the S-QU condition served as an active control against which the L-Bi and the L-QU pulses were compared. This approach was grounded in converging evidence indicating that short unidirectional pulses do not robustly alter corticospinal excitability (CSE) compared with longer pulses (Agboada et al., 2026). For instance, Halawa et al. (2021) showed that 120-μs and 100-μs L-Bi pulses delivered at 5 Hz increased CSE, whereas 80-μs pulses did not, suggesting that pulses must be longer to alter excitability. Similarly, concurrent rTMS-EEG evidence indicates that longer pulse durations (80-μs) increase local mean field power, whereas shorter pulses (30 μs) do not yield comparable changes in cortical excitability (Casula et al., 2018). Finally, McNaughton et al. (1978) found that long-term potentiation occurred only when pulse duration exceeded 120 μs during stimulation of the hippocampal formation. Taken together, these findings suggest that S-QU pulses provide a valid active control condition for isolating the effects of longer pulses on CSE. We hypothesized that unidirectional pulses would produce a larger increase in excitability than L-Bi pulses, and that L-QU pulses would be more effective than the S-QU, such that the L-QU condition would produce the strongest and most durable facilitatory aftereffects (Halawa et al., 2019; Wendt et al., 2023).

## Materials and Methods

### Participants and Procedure

Data were obtained from 17 right-handed, neurologically healthy volunteers aged 18 to 35 (11 females; mean age = 25.47, *SD* = 4.78). Recruitment was conducted through a public announcement on two university campuses in Sherbrooke, Canada. The study employed a single-blind, randomized, within-subject repeated-measures design consisting of three iTBS conditions and eight post-stimulation follow-up assessments at 10-minute intervals. Each participant attended three experimental visits at least 1 week apart, all held at the same time of day (Figure 1). The experiment took place at the *Centre Hospitalier Universitaire de Sherbrooke* (Quebec, Canada). Participants were compensated for their time with $50 CAN per visit. Two participants attended only one visit, and a third participant attended two of three visits. The study protocol was approved by the *Comité d’éthique de la recherche du CIUSS de l’Estrie – Centre Hospitalier Universitaire de Sherbrooke*, in accordance with the Declaration of Helsinki. Written informed consent was obtained from all participants after confirming their eligibility to undergo the TMS protocol using a standardized questionnaire, in accordance with the most recent TMS procedural safety guidelines (Rossi et al., 2021).

**Figure 1.**
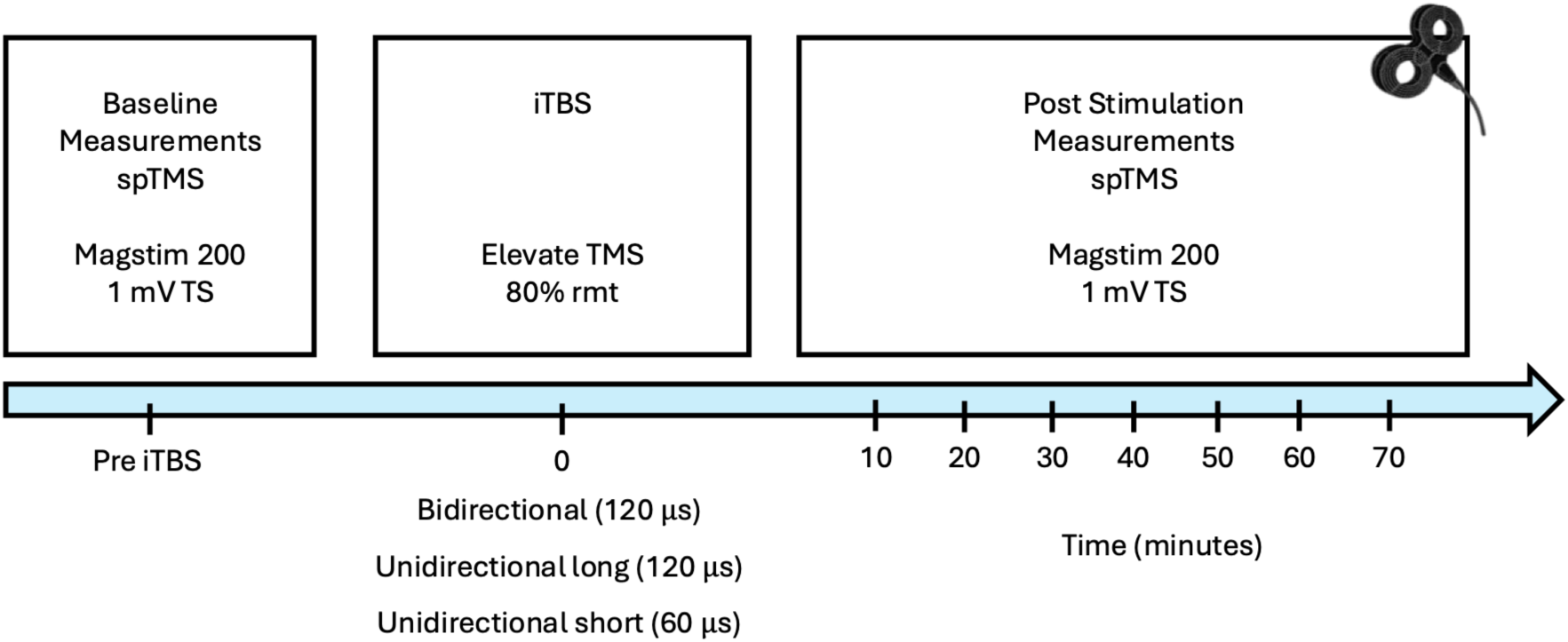
Protocol Overview.

### TMS Stimulation and MEP Recording

Single pulses were delivered over the left primary motor cortex (M1) using a Magstim 200 device (Magstim, Whitland, UK) with a 70-mm figure-of-eight coil in the anterior-to-posterior (AP) direction and a unidirectional pulse shape. The coil was placed at a 45° angle from the midline of the head. iTBS was administered using a cTMS apparatus with a 70-mm-diameter figure-of-eight air-cooled coil (Elevate TMS, Rogue Research, Canada).

At the start of the experiment, single-pulse TMS and neuronavigation were used to localize the motor hotspot – the area of the primary motor cortex that reliably produces large MEPs in the contralateral hand. We then determined the resting motor threshold (RMT) – the minimal stimulation intensity that elicits MEPs with a peak-to-peak amplitude of at least 50 μV in 5 of 10 trials. Next, we set the test-stimulus (TS) intensity, the baseline level of excitability, starting at 120% of the RMT and adjusting by 2-3% to obtain an average MEP amplitude of 1 mV in the contralateral right first dorsal interosseus (Rossini et al., 2015). Twenty-one MEPs were acquired at each acquisition time point: one before iTBS and 7 times after, starting at 10 minutes post-stimulation, with follow-up measures every 10 minutes up to 70 minutes. This number of trials has been shown to give a reliable measure of corticospinal excitability (Chang et al., 2016).

iTBS was administered using a cTMS apparatus (Elevate TMS, Rogue Research, Montreal, Canada) applied to the motor hotspot. iTBS consisted of triplets of pulses delivered at 50 Hz, with triplets repeated every 200 ms (5 Hz) for 2 s, followed by an 8-s pause. Stimulation was set at 80% of the RMT on the cTMS, established prior to the administration with the pulse parameters of interest. Across the three visits, we manipulated two pulse parameters: (1) the M-ratio (the relative amplitude of the second phase with respect to the first phase of the induced electric field pulse), and (2) the first-phase pulse width. The three conditions were: (1) long bidirectional (L-Bi) (M-ratio = 0.690, first-phase width=120µs), (2) long quasi-unidirectional (L-QU) (M-ratio = 0.370, first-phase width=120µs), and (3) control (see explanation above) short quasi-unidirectional (S-QU) (M-ratio = 0.370, first-phase width=60µs). The corresponding second-phase durations were 280 µs for both unidirectional conditions and 200 µs for the L-Bi condition (Figure 2). Thus, the S-QU and L-QU conditions differed in first-phase pulse width while maintaining the same M-ratio, whereas the long quasi-unidirectional and L-Bi conditions had the same first-phase duration.

**Figure 2.**
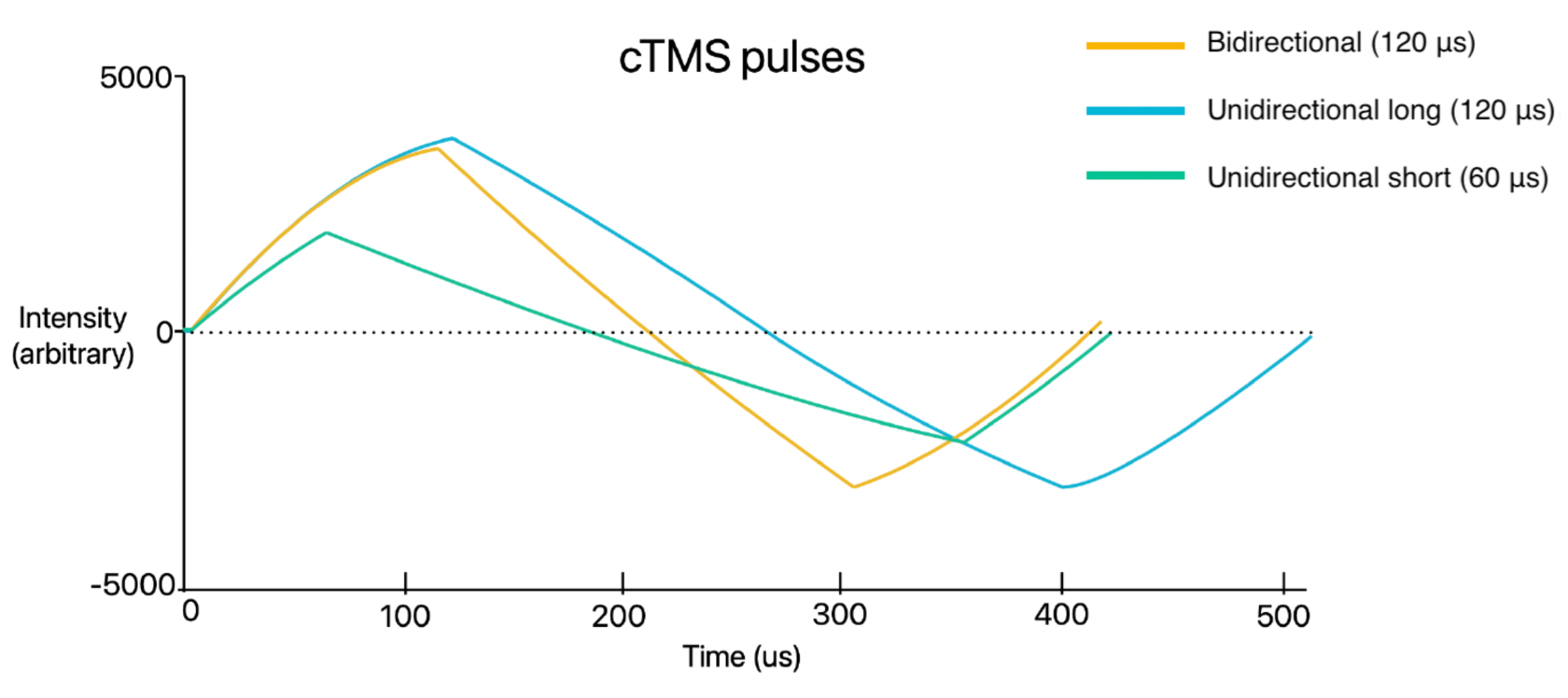
iTBS pulses as depicted on the cTMS. The 3 pulse types are depicted on the same intensity scale over time. The bidirectional pulse (M-ratio =0.690) has a positive phase of 120 µs and a negative phase of 200 µs. The L-QU pulse (M-ratio =0.370) has a positive phase of 120 µs and a negative phase of 280 µs. The S-QU pulse (M-ratio =0.370) has a positive phase lasting 60 µs and a negative phase lasting 280 µs.

Neuronavigation was used during each visit to ensure stable coil positioning during CSE measurement and iTBS administration (BrainSight, Rogue Research, Canada). LabChart 7 (ADI Instruments, Colorado Springs, USA) was used for electromyography (EMG) recording at a sampling rate of 4 kHz. The low-pass filter was filtering at 0.3 Hz and the high-pass filter at 1 kHz. All data were stored offline for analysis. Electrodes were placed in a belly-tendon arrangement with a ground electrode placed on the right elbow. Throughout the visit, participants’ hands remained at rest on a pillow, in supination. There was no serious adverse event to report.

### Data processing and analyses

Before processing, MEPs were assessed for the presence of outliers within each acquisition time point and removed before model fitting using a single round of median absolute deviation (MAD) with a cutoff of 3 MADs (Leys et al., 2013). This resulted in the rejection of 4.58% of trials and was performed to ensure that excitability estimates would not be skewed by outliers. The remaining MEPs were averaged separately for each time point, condition, and participant, and ratios were computed using the baseline measurement as the denominator for each time point.

A mixed linear model (MLM) was fit using restricted maximum likelihood with the GAMLj3 module in Jamovi (v. 2.7.13). Mixed linear models are a strong choice for physiological studies because they account for within- and between-group variability in the estimates and the non-independence of the data (Harrison et al., 2018). The fixed effects were time points (baseline and 10-70 minutes post-stimulation), pulse parameters (L-Bi, L-QU, S-QU), and their interaction. The random-effects structure included intercepts and allowed for participant-level variation in response to iTBS. For analysis, we retained the higher-order model that converged, was nonsingular, and yielded the lowest relative Akaike (AIC) value (Harrison et al., 2018).

To decompose interactions, we performed simple-effects analyses for each condition across all time points to assess whether CSE differed from baseline after iTBS. We then performed pairwise post hoc comparisons. We applied the Benjamini-Hochberg (BH) correction to control the false discovery rate (q = .05) (Benjamini & Hochberg, 1995).

## Results

### cTMS Motor Thresholds

A mixed linear model revealed a main effect of pulse parameters on RMTs (*F* (2, 27.05) = 13.96, *p* < .001); RMT was lowest for the L-Bi (28% ± 4.50), followed by the L-QU (32% ± 4.84) and the S-QU (36% ± 6.32). Post hoc pairwise comparisons indicated significantly lower RMTs for the L-Bi condition compared with the L-QU condition (*t* (27.2) = −2.74, *p* = .011), and the S-QU condition (*t* (27) = −6.40, *p* < .001). RMT was also significantly lower for the L-QU compared with the S-QU condition (*t* (27) = −2.76, *p* = .011). Thus, RMT increased progressively from the L-Bi to the L-QU and S-QU conditions (Figure 3).

**Figure 3.**
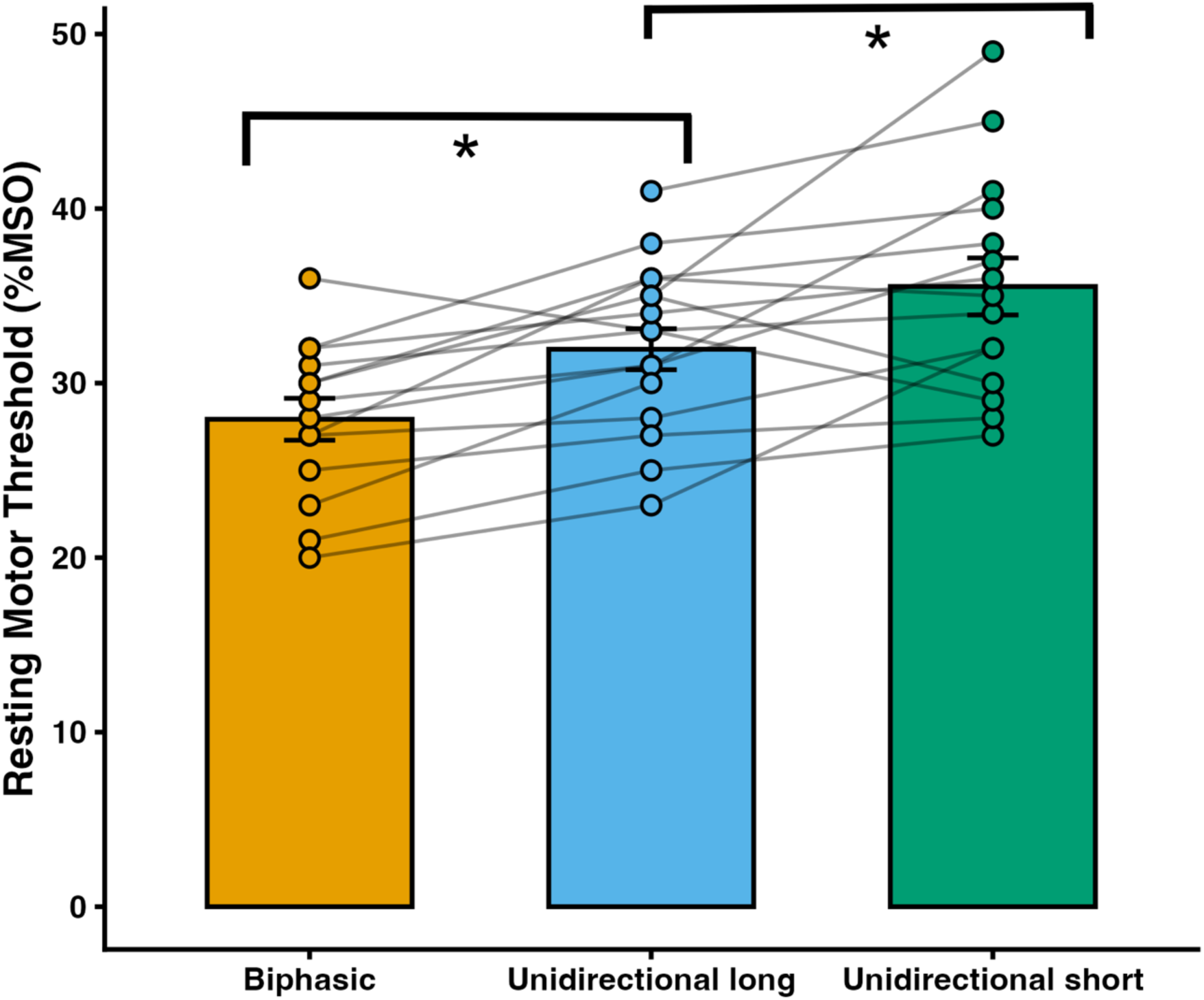
Resting Motor Thresholds Obtained on the cTMS. The bidirectional pulse lowered the motor threshold the most on the cTMS. The L-Bi motor thresholds are, on average 28% (SD= 4.50), L-QU, 32% (SD= 4.84) and S-QU, 36% (SD= 6.32). The error bars represent the between-subjects standard error of the mean (SEM).

### Corticospinal Excitability Changes induced by iTBS

The omnibus test of the fixed effects of the linear model revealed a significant main effect of time (χ2 (7) = 19.27, *p* <.001), condition (χ2 (2) = 105,54, *p* <.001), and interaction (χ2 (14) = 3.67, *p* <.001), indicating that the increase in cortical excitability depended on both time and pulse parameters used. Post hoc analyses showed significant differences between all conditions, where the L-Bi induced a larger increase in amplitude than the L-QU (*t* (7163) = 24.8, *p* < .001) and the S-QU (*t* (7151) = 36.4, *p* < .001), and the L-QU more than the S-QU (*t* (7163) = 11.6, *p* < .001; Figure 4B)

**Figure 4.**
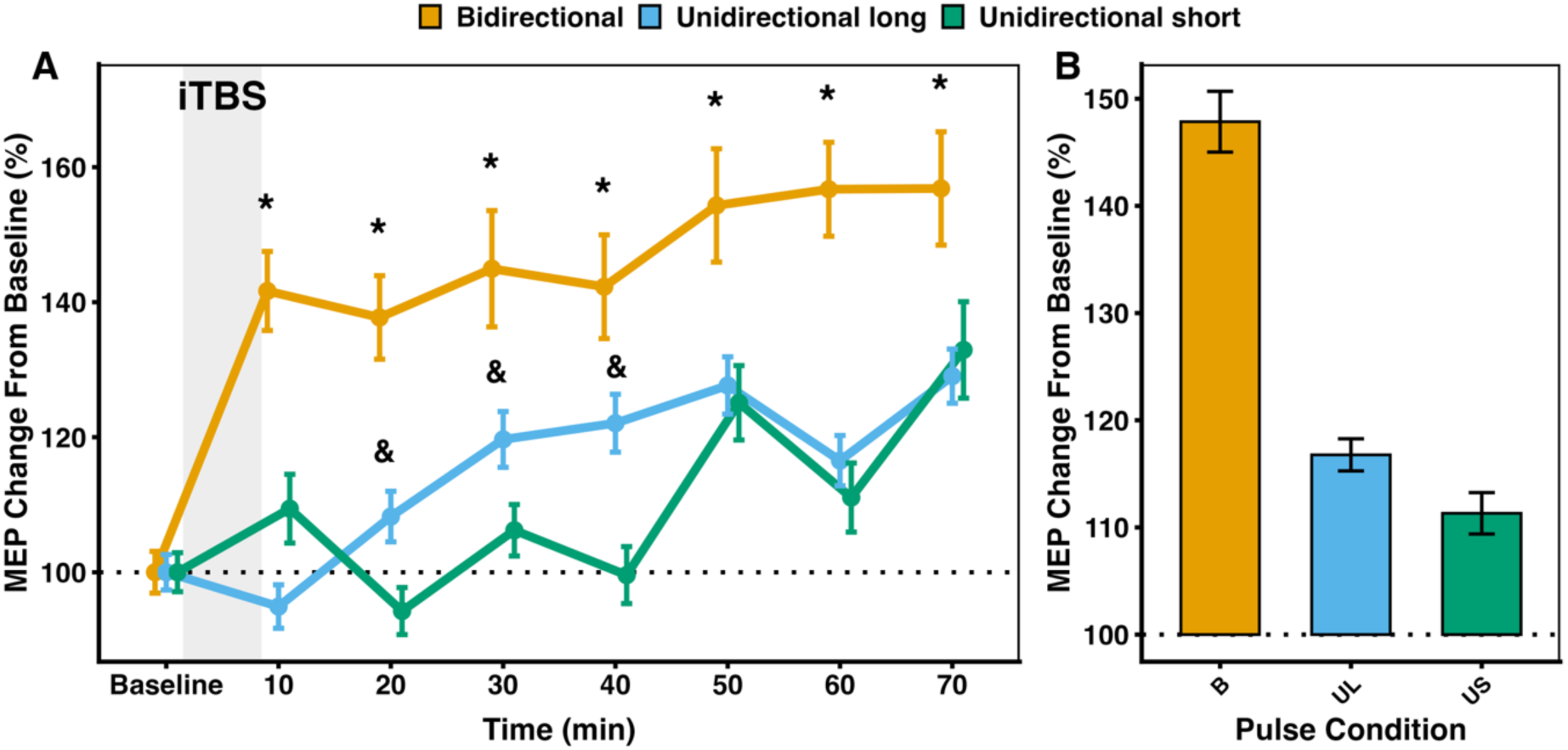
**(A)** Estimated Marginal Means of the MEP amplitude changes from baseline. The L-Bi pulses increased MEP amplitudes more than both unidirectional pulses at all time points (*). The L-QU pulse increased MEPs more than the S-QU pulses at times 20 and 30 (&). (B) Estimated marginal means of MEP amplitude change from baseline by condition, converging across all time points. The error bars represent the between-subjects standard error of the mean (SEM).

Simple effects of pulse parameters across time points show differences at all time points, except for baseline. The L-Bi pulse increased excitability significantly more than the L-QU and S-QU pulses at all time points. Specifically, significant differences are present at 10 minutes post iTBS (both *p* < .001), 20 minutes (both *p* < .001), 30 minutes (Long; p = .003, Short; p < .001), 40 minutes (Long; p = .014, Short; p < .001), 50 minutes (both *p* < .001), 60 minutes (both *p* < .001), and 70 minutes (both *p* < .001). Moreover, the L-QU pulse induced more excitability than the S-QU pulse at 20 minutes (*p* = .008), 30 minutes (*p* = .008) and 40 minutes (*p* = .002) after iTBS (Figure 4A).

We then ran simple effects of time points across pulse parameters to examine whether each condition significantly changed CSE relative to baseline. After BH correction, we found that the L-Bi pulse increased CSE significantly from baseline across all post time points (all *p* < .001), the L-QU pulse from 30 to 70 minutes after iTBS (all *p* < .05), and the S-QU pulse at 50 and 70 minutes after iTBS (both *p* < .001).

### Responders versus non-responders’ analysis

To determine which subjects responded to each stimulation with significant changes in CSE, the standard error of the mean (SEM) was calculated for each participant’s baseline trials in the three conditions. Then, the grand average SEM of the participants’ visits was multiplied by 1.96, the z-score threshold for Cronbach’s alpha at .05, corresponding to a 5% tolerance for error. For subjects who had facilitation of CSE the product was added to one, and the product of those who responded with inhibition was subtracted from one. If the final percentage change from baseline exceeded the average SEM, there was a 95% chance that the results indicated a significant change in excitability, leading to the classification of participants as responders, non-responders, or paradoxical responders. Most individuals responded to more than one condition (*n* = 10). Four participants responded to all conditions, and three participants responded to 1 condition only (L-Bi, *n* = 2; L-QU, *n* = 1). Paradoxical responses were distributed equally across conditions. (Figure 5).

**Figure 5.**
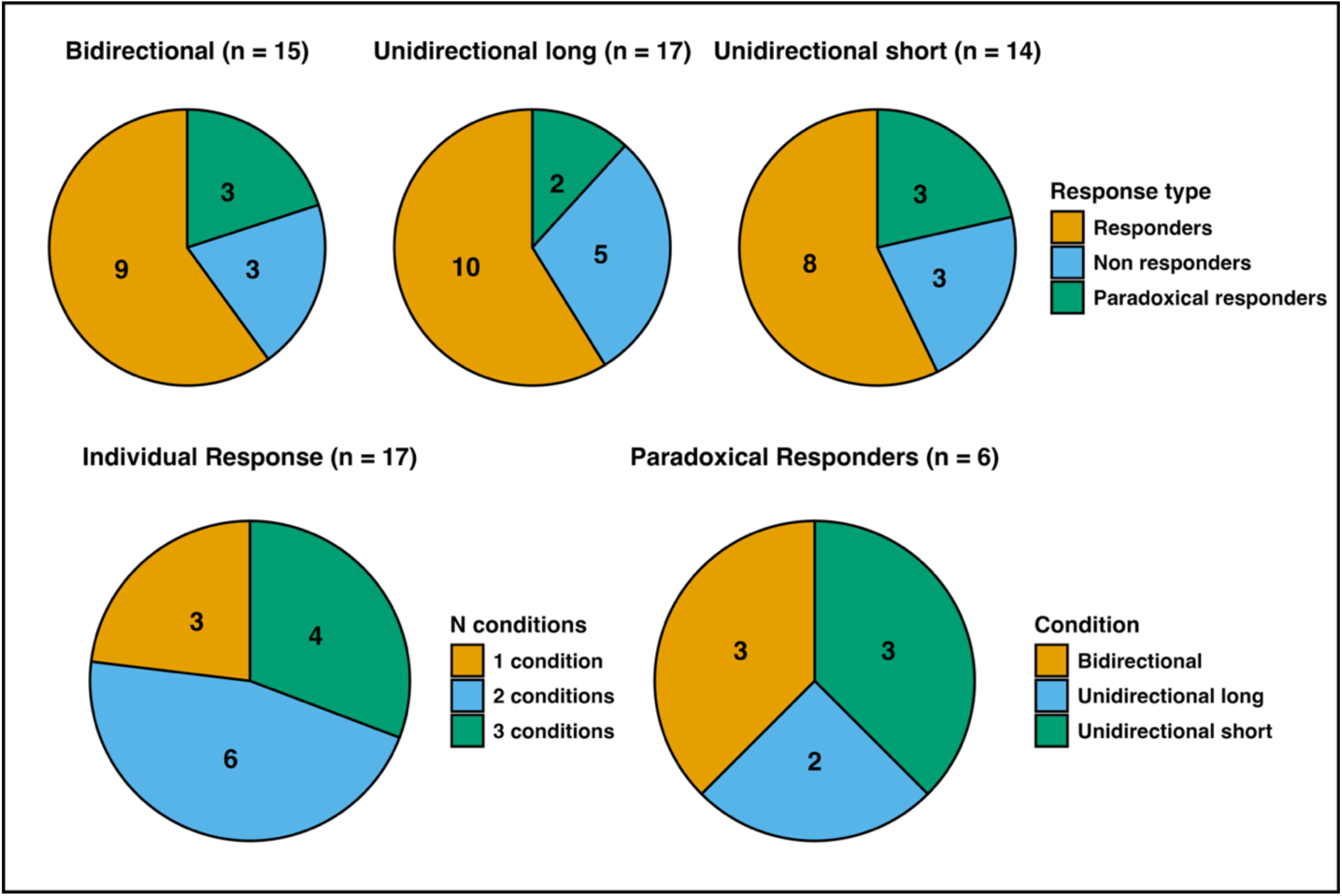
Responders to iTBS.

## Discussion

This study examined the effects of three active iTBS conditions that differed in pulse shape and width on CSE. Contrary to the initial hypothesis, the L-QU pulse did not induce the greatest increase in CSE. Instead, the L-Bi condition induced the largest and most sustained facilitation, greater than the unidirectional conditions from 10 to 70min after iTBS. The L-QU condition also increased CSE, but the effect was smaller and appeared later, whereas the S-QU condition produced the weakest effects, with significant changes only at 50 and 70 min. Thus, although the magnitude of facilitation differed between conditions, all three seemed to produce lasting physiological effects. Furthermore, the distinct time courses observed suggest that pulse parameters may influence not only the extent of facilitation, but also the timing and duration of the iTBS effects on CSE (Wischnewski & Schutter, 2015), highlighting the importance of using multiple time points when comparing stimulation protocols aiming at modulating CSE.

The stronger facilitation observed in the L-Bi condition is in contrast with findings from previous studies using conventional rTMS. Namely, 1 Hz (Halawa et al., 2019) and 5 Hz (Halawa et al., 2021) rTMS protocols using unidirectional pulses induced respectively stronger inhibition and facilitation than conditions with biphasic pulses. Similarly, Wendt and colleagues (2023) reported greater facilitation following iTBS using unidirectional pulses in comparison to biphasic pulses. However, it should be noted that while these studies compared unidirectional and bidirectional pulses, they differed with regards to pulse durations, pulse width, and TMS apparatus, all of which may have contributed to the discrepant findings.

One possible explanation for the greater facilitation resulting from the L-Bi condition is that bidirectional pulses recruit a broader population of cortical neurons than unidirectional pulses. Indeed, unidirectional pulses are believed to preferentially activate neurons sensitive to current flowing in one direction, whereas the second phase of a bidirectional pulse may recruit an additional population with a different directional sensitivity (Sommer et al., 2018). During iTBS, the recruitment of multiple neuronal populations by the L-Bi could increase the pool of synapses available for the cumulative, burst-dependent processes underlying LTP-like plasticity. Conversely, the greater selectivity of unidirectional stimulation may restrict recruitment to a narrower neuronal population, potentially limiting synaptic summation (Sommer et al., 2018). Bidirectional pulses may also produce steeper membrane polarization during their second phase in response to initial hyperpolarization, which could contribute to their greater facilitatory potential (Delvendahl et al., 2014). Near-rectangular bidirectional pulses may be particularly well suited to the repeated-burst structure of iTBS. However, this interpretation remains speculative because evidence regarding the effects of pulse directionality and waveform on neuronal recruitment has largely been derived from single-pulse studies (Agboada et al., 2026). Further work directly examining neuronal recruitment during iTBS will be necessary to determine the mechanisms underlying the greater facilitation observed with L-Bi stimulation.

Pulse shape may also contribute to the observed differences between conditions. Sinusoidal and near-rectangular pulses have been shown to produce distinct neurophysiological effects (Peterchev et al., 2014; Agboada et al., 2026). For example, unidirectional sinusoidal pulses have been reported to produce greater facilitation than bidirectional pulses following 10-Hz rTMS (Arai et al., 2007), whereas Goetz et al. (2016) found greater inhibitory after-effects with unidirectional rectangular pulses than with other pulse shapes during 1-Hz rTMS. However, direct comparisons of near-rectangular and sinusoidal pulses under otherwise equivalent bidirectional and unidirectional iTBS conditions are lacking. Such studies are needed to determine whether the enhanced facilitation observed with L-Bi in the present study is attributable primarily to pulse directionality, pulse shape, or their interaction.

Increasing the first-phase pulse width from 60 to 120 µs also reduced resting motor threshold (RMT) and enhanced the magnitude of iTBS after-effects. The reduction in RMT is consistent with the strength–duration relationship, whereby longer pulses require lower stimulation intensities to activate cortical neurons (Peterchev et al., 2013). Previous work has similarly shown that increasing pulse width can lower motor threshold and MEP latency while increasing MEP amplitude (D’Ostilio et al., 2016; Halawa et al., 2019). The greater effectiveness of longer pulses may reflect the recruitment of neuronal populations with longer membrane time constants, which would stay tuned for longer durations (Stern et al., 2015; Halawa et al., 2019). Thus, pulse width represents a potentially important, yet relatively underexamined, parameter for optimizing high-frequency TMS protocols.

Despite the differences in mean facilitation across conditions, responder analysis did not identify a single pulse condition associated with a greater likelihood of a significant facilitatory response. Although, most participants responded to more than one condition, this overlap was far from ubiquitous, indicating substantial inter-individual variability. Paradoxical responses to iTBS were also observed across conditions rather than being associated with a specific pulse configuration. Individual neurophysiological characteristics may contribute to the direction and magnitude of the response to iTBS. For example, individuals with shorter baseline MEP latencies have been reported to show greater CSE facilitation following continuous TBS, potentially reflecting differences in the neuronal populations recruited by stimulation (Huang & Mouraux, 2015). More broadly, baseline cortical excitability and time-varying physiological states of M1 may influence responsiveness to TBS. Characterizing individuals’ baseline neurophysiology in future studies may therefore help explain the substantial within- and between-individual variability observed in response to different TBS protocols.

### Limitations and future directions

Several limitations should be considered. First, the sample was small, and some participants did not complete all three visits. Second, the study did not include a sham condition; thus, we cannot completely exclude that CSE would be modulated by time alone throughout the experimental session alone, even if this is unlikely. Third, the design was not fully factorial; as a result, in the absence of a short bidirectional condition, the independent effects of pulse width and M-ratio cannot be determined. Also, the L-Bi and L-QU conditions differed in second-phase width, which will require further investigation. Considering that the iTBS intervention and MEP measurements were conducted using different stimulators and coils, it is possible that the circuits stimulated with unidirectional single pulses during CSE measurements may not be the same as those engaged during the iTBS procedure, particularly in the bidirectional iTBS condition. However, this appears not to be an issue, as iTBS with bidirectional pulses led to more CSE increase, even when assessed with unidirectional single pulses. Future studies comparing conventional and near-rectangular waveforms using the same stimulation platform could also help explain the differences between the present findings and those of Wendt et al. (2023). Lastly, one should be cautious regarding the extrapolation of the present results to cortical regions outside of M1, as its cytoarchitecture is particular and different from the one seen in prefrontal regions, where iTBS is often applied for clinical applications. In this regard, conducting TMS-EEG studies could clarify if these results can be replicated in other brain regions.

## Conclusion

Here, we showed that iTBS administered using 120µs-width L-Bi pulses induced a larger and more sustained increase in CSE in comparison to other unidirectional pulses. Of interest, the L-QU pulse was also more effective than S-QU pulses, supporting pulse width as an important parameter for high-frequency rTMS. We also noted significant variability with regards to the aftereffects of iTBS. The present findings suggest that pulse shape and pulse width both contribute to the aftereffects of iTBS, opening the door to tailoring pulse parameters to optimize response to rTMS intervention, and possibly improve response and remission rates in clinical settings.

## Acknowledgements

This work was funded by the Natural Sciences and Engineering Research Council of Canada (NSERC), the Fondation Enfant-Soleil, and the Fonds de Recherche du Québec – Santé (FRQS) awarded to JFL. BP is supported by the Molson Foundation and AJF is supported by the Canadian Institutes of Health Research (CIHR). ST is supported by an award from the FRQS.

## AI statement

The use of AI in this work was limited to checking grammar and spelling errors.

## References

1. Agboada, D., Rethwilm, R., Kuder, M., Mack, W., & Seiberl, W. (2026). A systematic review of the effect of pulse parameters of next-generation TMS devices on corticospinal excitability and neuroplasticity. Brain Research, 1873, 150120. 10.1016/j.brainres.2025.150120

2. Arai, N., Okabe, S., Furubayashi, T., Mochizuki, H., Iwata, N. K., Hanajima, R., Terao, Y., & Ugawa, Y. (2007). Differences in after-effect between monophasic and biphasic high-frequency rTMS of the human motor cortex. Clinical Neurophysiology, 118(*10*), 2227–2233. 10.1016/j.clinph.2007.07.006

3. Benjamini, Y., & Hochberg, Y. (1995). Controlling the False Discovery Rate: A Practical and Powerful Approach to Multiple Testing. Journal of the Royal Statistical Society. Series B (Methodological), 57(*1*), 289–300. http://www.jstor.org/stable/2346101

4. Blumberger, D. M., Vila-Rodriguez, F., Thorpe, K. E., Feffer, K., Noda, Y., Giacobbe, P., Knyahnytska, Y., Kennedy, S. H., Lam, R. W., Daskalakis, Z. J., & Downar, J. (2018). Effectiveness of theta burst versus high-frequency repetitive transcranial magnetic stimulation in patients with depression (Three-d) : A randomised non-inferiority trial. The Lancet, 391(10131), 1683–1692. 10.1016/S0140-6736(18)30295-2

5. Casula, E. P., Rocchi, L., Hannah, R., & Rothwell, J. C. (2018). Effects of pulse width, waveform and current direction in the cortex : A combined cTMS-EEG study. Brain Stimulation, 11(5), 1063–1070. 10.1016/j.brs.2018.04.015

6. Chang, W. H., Fried, P. J., Saxena, S., Jannati, A., Gomes-Osman, J., Kim, Y.-H., & Pascual-Leone, A. (2016). Optimal number of pulses as outcome measures of neuronavigated transcranial magnetic stimulation. Clinical Neurophysiology: Official Journal of the International Federation of Clinical Neurophysiology, 127(*8*), 2892–2897. 10.1016/j.clinph.2016.04.001

7. Chaves, A. R., Cuda, J., Shim, S., Drodge, J., Nasr, Y., Brender, R., Antochi, R., McMurray, L., Pilutti, L. A., & Tremblay, S. (2024). Optimizing antidepressant benefits : Effect of theta burst stimulation treatment in physically active people with treatment-resistant depression. Journal of Affective Disorders, 367, 876–885. 10.1016/j.jad.2024.09.024

8. Delvendahl, I., Lindemann, H., Jung, N. H., Pechmann, A., Siebner, H. R., & Mall, V. (2014). Influence of waveform and current direction on short-interval intracortical facilitation : A paired-pulse tms study. Brain Stimulation, 7(1), 49–58. 10.1016/j.brs.2013.08.002

9. D’Ostilio, K., Goetz, S. M., Hannah, R., Ciocca, M., Chieffo, R., Chen, J.-C. A., Peterchev, A. V., & Rothwell, J. C. (2016). Effect of coil orientation on strength– duration time constant and I-wave activation with controllable pulse parameter transcranial magnetic stimulation. Clinical Neurophysiology, 127(1), 675–683. 10.1016/j.clinph.2015.05.017

10. Goetz, S. M., Luber, B., Lisanby, S. H., Murphy, D. L. K., Kozyrkov, I. C., Grill, W. M., & Peterchev, A. V. (2016). Enhancement of neuromodulation with novel pulse shapes generated by controllable pulse parameter transcranial magnetic stimulation. Brain Stimulation, 9(1), 39–47. 10.1016/j.brs.2015.08.013

11. Halawa, I., Reichert, K., Aberra, A. S., Sommer, M., Peterchev, A. V., & Paulus, W. (2021). Effect of pulse duration and direction on plasticity induced by 5 Hz repetitive transcranial magnetic stimulation in correlation with neuronal depolarization. Frontiers in Neuroscience, 15. 10.3389/fnins.2021.773792

12. Halawa, I., Shirota, Y., Neef, A., Sommer, M., & Paulus, W. (2019). Neuronal tuning : Selective targeting of neuronal populations via manipulation of pulse width and directionality. Brain Stimulation, 12(*5*), 1244–1252. 10.1016/j.brs.2019.04.012

13. Harrison, X. A., Donaldson, L., Correa-Cano, M. E., Evans, J., Fisher, D. N., Goodwin, C. E. D., Robinson, B. S., Hodgson, D. J., & Inger, R. (2018). A brief introduction to mixed effects modelling and multi-model inference in ecology. PeerJ, 6, e4794. 10.7717/peerj.4794

14. Huang, G., & Mouraux, A. (2015). Mep latencies predict the neuromodulatory effect of ctbs delivered to the ipsilateral and contralateral sensorimotor cortex. PLOS ONE, 10(8), e0133893. 10.1371/journal.pone.0133893

15. Huang, Y. Z., Edwards, M. J., Rounis, E., Bhatia, K. P., & Rothwell, J. C. (2005). Theta burst stimulation of the human motor cortex. Neuron, 45(*2*), 201–206. 10.1016/j.neuron.2004.12.033

16. Leys, C., Ley, C., Klein, O., Bernard, P., & Licata, L. (2013). Detecting outliers : Do not use standard deviation around the mean; use absolute deviation around the median. Journal of Experimental Social Psychology, 49(*4*), 764–766. 10.1016/j.jesp.2013.03.013

17. McNaughton, B. L., Douglas, R. M., & Goddard, G. V. (1978). Synaptic enhancement in fascia dentata : Cooperativity among coactive afferents. Brain Research, 157(2), 277–293. 10.1016/0006-8993(78)90030-6

18. Peterchev, A. V., Goetz, S. M., Westin, G. G., Luber, B., & Lisanby, S. H. (2013). Pulse width dependence of motor threshold and input–output curve characterized with controllable pulse parameter transcranial magnetic stimulation. Clinical Neurophysiology, 124(7), 1364–1372. 10.1016/j.clinph.2013.01.011

19. Peterchev, A. V., D’Ostilio, K., Rothwell, J. C., & Murphy, D. L. (2014). Controllable pulse parameter transcranial magnetic stimulator with enhanced circuit topology and pulse shaping. Journal of neural engineering, 11(*5*), 056023. 10.1088/1741-2560/11/5/056023

20. Rossi, S., Antal, A., Bestmann, S., Bikson, M., Brewer, C., Brockmöller, J., Carpenter, L. L., Cincotta, M., Chen, R., Daskalakis, J. D., Di Lazzaro, V., Fox, M. D., George, M. S., Gilbert, D., Kimiskidis, V. K., Koch, G., Ilmoniemi, R. J., Lefaucheur, J. P., Leocani, L., … Hallett, M. (2021). Safety and recommendations for TMS use in healthy subjects and patient populations, with updates on training, ethical and regulatory issues : Expert Guidelines. Clinical Neurophysiology, 132(*1*), 269–306. 10.1016/j.clinph.2020.10.003

21. Rossini, P. M., Burke, D., Chen, R., Cohen, L. G., Daskalakis, Z., Di Iorio, R., Di Lazzaro, V., Ferreri, F., Fitzgerald, P. B., George, M. S., Hallett, M., Lefaucheur, J. P., Langguth, B., Matsumoto, H., Miniussi, C., Nitsche, M. A., Pascual-Leone, A., Paulus, W., Rossi, S., … Ziemann, U. (2015). Non-invasive electrical and magnetic stimulation of the brain, spinal cord, roots and peripheral nerves : Basic principles and procedures for routine clinical and research application. An updated report from an I.F.C.N. Committee. Clinical Neurophysiology, 126(*6*), 1071–1107. 10.1016/j.clinph.2015.02.001

22. Sommer, M., Alfaro, A., Rummel, M., Speck, S., Lang, N., Tings, T., & Paulus, W. (2006). Half sine, monophasic and biphasic transcranial magnetic stimulation of the human motor cortex. Clinical Neurophysiology, 117(4), 838–844. 10.1016/j.clinph.2005.10.029

23. Sommer, M., Ciocca, M., Chieffo, R., Hammond, P., Neef, A., Paulus, W., Rothwell, J. C., & Hannah, R. (2018). TMS of primary motor cortex with a biphasic pulse activates two independent sets of excitable neurones. Brain Stimulation, 11(3), 558–565. 10.1016/j.brs.2018.01.001

24. Stern, S., Agudelo-Toro, A., Rotem, A., Moses, E., & Neef, A. (2015). Chronaxie measurements in patterned neuronal cultures from rat hippocampus. PLOS ONE, 10(*7*), e0132577. 10.1371/journal.pone.0132577

25. Wendt, K., Sorkhabi, M. M., Stagg, C. J., Fleming, M. K., Denison, T., & O’Shea, J. (2023). The effect of pulse shape in theta-burst stimulation : Monophasic vs biphasic TMS. Brain Stimulation, 16(*4*), 1178–1185. 10.1016/j.brs.2023.08.001

26. Wischnewski, M., & Schutter, D. J. L. G. (2015). Efficacy and time course of theta burst stimulation in healthy humans. Brain Stimulation, 8(4), 685–692. 10.1016/j.brs.2015.03.004

